# DeepSipred: A deep-learning-based approach on siRNA inhibition prediction

**DOI:** 10.1101/2023.11.02.565277

**Authors:** Bin Liu, Huiya Huang, Weixi Liao, Xiaoyong Pan, Cheng Jin, Ye Yuan

## Abstract

**Motivation:** The use of exogenous small interfering RNAs (siRNAs) for gene silencing has become a widespread molecular tool for gene function study and new drug identification. Although the pathway of RNAi to mediate gene expression has been widely investigated, the selection of hyperfunctional siRNA with high inhibition remains challenging.

**Results:** In this study, we build a deep-learning-based approach on siRNA inhibition prediction, named DeepSipred. It combines features from sequence context, thermodynamic property, and other expert knowledge together to predict the inhibition more accurately than existing methods. The sequence features from siRNA and local target mRNA are generated via one-hot encoding and pretrained RNA-FM encoding. The convolution layers with multiple kernels in DeepSipred can detect various decisive motifs, which will determine the actual inhibition of siRNA. The thermodynamic features are calculated from Gibbs Free Energy. In addition, the expert knowledge includes those design criteria from previous studies. Benchmarked on large available public datasets, the 10-fold cross-validation results indicate that our predictor achieving the state-of-the-art performance.

## 1 Introduction

RNA interference (RNAi) is common and spontaneous in many eukaryotes as a cellular pathway in posttranscriptional gene regulation. RNAi-based knockdown of a specific gene has been a widely used molecular technology in recent years [1]. One of the main regulators is 21∼23-nucleotide small interfering RNA (siRNA), which is first cleaved from a long double-stranded RNA (dsRNA) by the ribonuclease-III enzyme Dicer2, and then the antisense strand (AS) (also called the guide strand) in the siRNA duplex is assembled into the Argonaute protein and the catalytic media of the RNA-induced silencing complex (RISC). RISC can cleave target mRNAs that hybridize with the AS by complementary Watson-Crick base pairing to induce degradation or repress the posttranslational process by partial complementarity between the seed region of the AS and the 3’ untranslated regions (3’ UTRs) of target RNAs [2] [3] [4] [5]. SiRNAs have gradually played an important role in investigating gene function and show great potential in therapeutic applications against cancers [6], HIV, polyglutanmine-induced neurodegeneration [7], dominantly inherited brain diseases, skin diseases [8], and others.

However, siRNA needs to be designed carefully to ensure its inhibition toward a target mRNA [9] [10]. Silencing inhibition is mainly determined by the sequence patterns, binding affinity, and secondary structure around the binding regions. Various statistical and computational algorithms have been developed to design and select the most effective and functional siRNA for the target mRNA.

Early approaches were mainly based on two principles to automatically design siRNAs. One is to utilize expert knowledge to build empirical functionality criteria, involving GC content, thermodynamic properties, secondary structure, and other properties [11] [12] [13] [14]. The other is to identify the nucleotide preferences for particular positions by combining cellular assays and statistical analysis on experimentally validated hyper-functional siRNAs [15]. However, both of them are summarized from rather small datasets with biases due to preselection based on some prior empirical criteria. The results on a larger collection of publicly available datasets show that these methods suffer from poor generalization and robustness in unseen scenarios, almost close to random classification [16].

Fortunately, an increasing number of experimentally validated siRNAs have been collected. Computational methods based on feature engineering are widely applied in siRNA inhibition prediction. Huesken *et al*. collected and published a rather large dataset composed of 2431 randomly selected siRNAs targeting 34 mRNAs. They developed a model named BIOPREDsi with the Stuttgart Neural Net Simulator to achieve a high Pearson correlation coefficient (PCC) of 0.66 on an independent test set [17]. Vert *et al*. designed an accurate and interpretable LASSO linear regression model called DSIR using two simple sets of sequence features, nucleotide preferences at particular positions and short asymmetric motifs, to achieve comparative accuracy as BIOPREDsi and comprehensively interpret the impacts of various sequence features [18]. However, these methods also suffer from incomplete, redundant, or biased information in their features.

Recently, convolutional neural networks (CNNs) have been successfully applied to siRNA analysis. Han *et al*. designed a CNN-based model to predict siRNA potency in a data-driven way, making use of the nonlinear mapping ability of convolution and activation function to detect potential patterns from two modalities: local mRNA sequence context and thermodynamic property of the AS, which might not be found or verified in prior studies [19]. However, this model only takes limited features into a rather simple network with only one hidden layer, restricting its performance. Nevertheless, this model achieves considerable improvement in PCC and the area under the receiver operating characteristic curve (AUC) compared with traditional methods.

Apart from CNNs, the graph neural network (GNN) is also a new computational algorithm to model biological entities and their relationships, with nodes signifying molecules and edges representing interactions [20]. Massimo *et al*. developed a GNN-based network for siRNA-mRNA interaction prediction for the first time. They calculated the 3-mer frequencies (64 features) as the property of the siRNA node, 4-mer frequencies (256 features) as the property of the mRNA node, and the thermodynamic parameters generated from Gibbs energy and the RNAup program as the property of the siRNA-mRNA-interaction node. They consider the inhibition as the label of the siRNA-mRNA-interaction node, achieving a better result than the CNN-based method. However, this graph is trained and tested via deductive reasoning, and it cannot predict the inhibition of new siRNA that is not included in a known dataset. Moreover, enriching the node features may further enhance its accuracy [21].

In this study, we first collected available public datasets and then constructed a novel deep learning-based model for siRNA inhibition prediction, named DeepSipred. It enriches the characteristics of sequence context via one-hot encoding and pretrained RNA foundation model (RNA-FM) [22]. Features also consist of thermodynamic properties, the secondary structure, the nucleotide composition, and other expert knowledge. Then, it utilizes different kernels to detect potential motifs in sequence embedding, followed by a pooling operation. Next, we concatenate the output of pooling and all other features together. After being normalized in batches, it is fed into a deep and wide network with a sigmoid activation function. The 10-fold cross-validation results show satisfying results of our model, achieving state-of-the-art performance on the main metrics.

## 2. Materials and Methods

### 2.1 Benchmark datasets

In this study, we collected 3,536 experimentally validated siRNAs with corresponding potency, as suggested in the work of Han *et al*. [23]. They come from the studies of Huesken [17], Reynolds [24], Vickers [12], Haborth [25], Takayuki [26], and Ui-Tei [27]. Since the datasets from Reynolds, Vickers, Haborth, and Ui-Tei are all rather small, we gather them together and name them dataset-RVHU. The dataset from Takayuki is the only complete dataset targeting all positions of a single mRNA (EGFP), and we name it dataset-T. The dataset from Huesken is abbreviated as dataset-H. The details of these datasets are summarized in **Table 1**. Some approaches during comparison, such as DSIR, lack source codes and are hard to reproduce, so we utilize the i-score website to obtain their inhibition predictions. However, the first two siRNAs in Dataset-T are discarded on this website. Therefore, we removed 2 siRNAs from Dataset-T here. In addition, two siRNAs in dataset-RVHU failed to find target-binding sites on reported mRNAs, so we discarded them.

**Table 1.** The details of benchmark datasets. Two siRNAs are removed in dataset-T due to the limitation of i-score website, and two are discarded in dataset-RVHU as a result of lacking target sites.

| Dataset | Num of siRNAs | Num of target mRNAs |
| --- | --- | --- |
| Dataset-H | 2431 | 34 |
| Dataset-RVHU | 405(407) | 11 |
| Dataset-T | 700(702) | 1 |

The inhibitions of dataset-H, dataset-RVHU, and dataset-T range from 0 to 134.1, -27.8 to 98.9, and 0 to 97, respectively. After normalizing them individually to the range of 0∼1, we sorted them in an inhibition-descending order. As shown in **Fig. 1**, the normalized inhibitions of all three datasets show a nearly normal probability distribution, where the average is 0.543, and the standard deviation is 0.187.

**Fig. 1:**
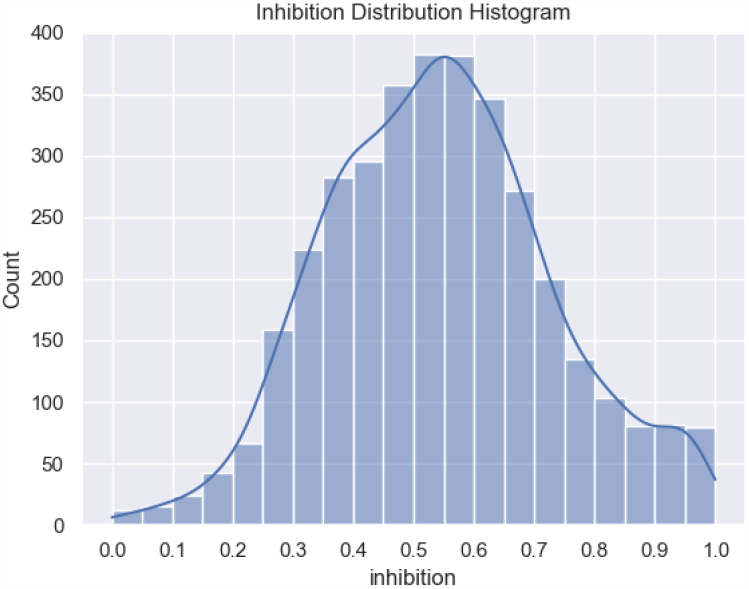
The distribution of the normalized inhibition. The average inhibition of 3536 siRNAs is 0.543, and the standard deviation is 0.187.

### 2.2 The siRNA inhibition predictor

The deep learning-based siRNA inhibition predictor consists of three modules (shown in **Fig. 2**): feature extraction, convolution and pooling, and full connection.

**Fig. 2:**
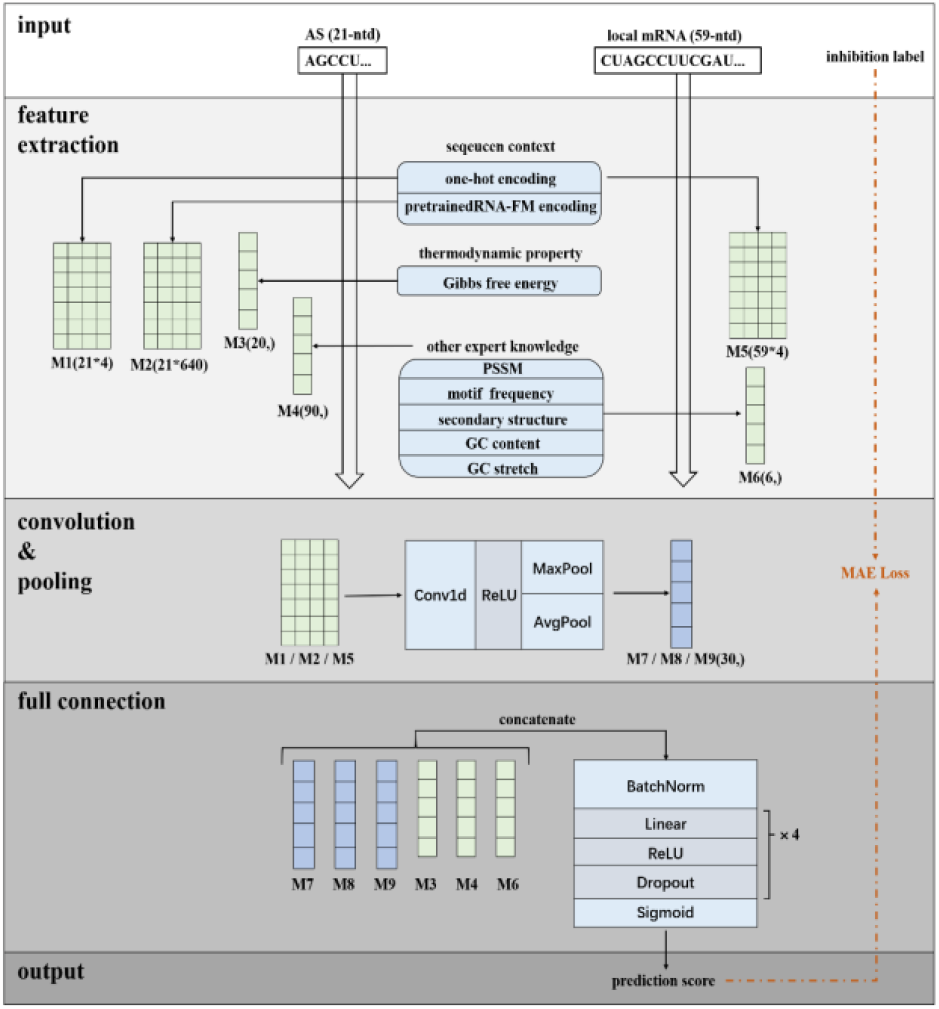
The architecture of our hybrid-deep-learning-based predictor for siRNA inhibition. The predictor first extracts features of siRNA and local mRNA sequence context via one-hot and pretrained RNA-FM encoding. The embeddings (M1, M2, and M5) are then processed by multiple one-dimensional convolutions with different kernels to detect potential motifs, followed by pooling operation. Then, the pooling results (M7, M8, and M9) are concatenated with other features, including thermodynamic property, the secondary structure, the nucleotide composition and so on (M3, M4, and M6). After batch normalization, a four-layer full connection module is used to aggregate information among diverse features. Finally, the model output a prediction score, ranging from 0 to 1.

#### 2.2.1 Feature Extraction Module

Here, we mainly consider three types of features, which come from sequence context, thermodynamic property, and some other prior expert knowledge.

##### 1) Encoding of siRNA and local mRNA sequence

A relevant study has shown that the flanking regions around the binding site on mRNA affect siRNA potency [28]. Here, we trim 20 nucleotides from the upstream and downstream regions on the target mRNA separately to form the local mRNA sequence, as suggested in previous work [23]. Note that the binding part usually refers to the core region of AS, composed of 19 nucleotides from its 5’ end. Therefore, the length of the local target mRNA adds up to 59 (20+19+20). We use one-hot encoding and pretrained RNA-FM encoding methods here. RNA-FM is a self-supervised model trained on 23 million ncRNA sequences from the RNAcentral database, and the produced embedding contains functional, sequential, structural, and evolutionary information [22]. Thus, we applied pretrained RNA-FM to help encode the structural and functional information of the siRNA sequence. One single nucleotide is transformed to a four-dimensional discrete vector by one-hot encoding and a 640-dimensional contiguous vector via pretrained RNA-FM encoding.

Overall, 21-nucleotide AS is transformed into two feature matrixes with sizes of 21*4 and 21*640. To save the computational cost caused by RNA-FM, the 59-nucleotide local mRNA is only encoded into one matrix with a size of 59*4 using one-hot encoding. If the binding site is located at the end of the mRNA, the upstream or downstream regions would contain less than 20 nucleotides. In this case, we encode the missing nucleotide with a special vector of [0.05, 0.05, 0.05, 0.05].

##### 2) Encoding of siRNA thermodynamic properties

The thermodynamic stability of siRNA duplexes plays a vital role in the inhibition potency and longevity of siRNAs [26]. The absolute and relative thermodynamic stabilities of bases at the 5’ and 3’ ends of siRNAs determine which strand is bound to the RISC and hybridized with the target mRNA [2] [11] [29].

Here, we mainly consider the Gibbs free energy of the core region of AS. There are a total of 20 thermodynamic parameters, as shown in **Table 2**, including the energy of every two adjacent base pairs, the difference between the 5’ and 3’ ends, and the overall energy.

**Table 2.**
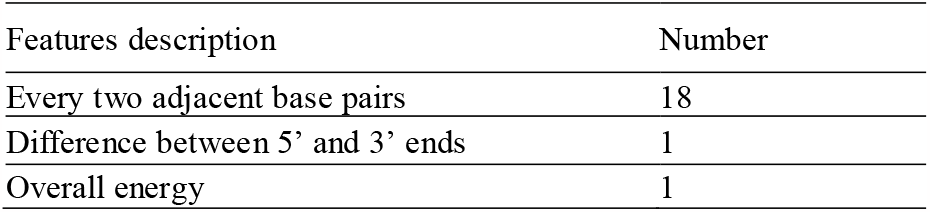
The description of thermodynamic property (Gibbs free energy) in the core region of AS.

| Features description | Number |
| --- | --- |
| Every two adjacent base pairs | 18 |
| Difference between 5' and 3' ends | 1 |
| Overall energy | 1 |

##### 3) Encoding of other biological features from expert knowledge

In addition to sequence context and thermodynamic properties, we extract four groups of features from expert knowledge.

###### Frequencies of multiple motifs

The sequence composition plays a primary role in determining the position of Dicer cleavage in both dsRNA and short hairpin RNA [30] [31]. Here, we calculate the occurrences of 4, 16, and 64 kinds of possible motifs in the 1-mer, 2-mer, and 3-mer counts of the siRNA sequence, respectively. This provides a simple way to encode the presence of specific nucleotides and short motifs.

###### Position Specific Scoring Matrix (PSSM)

It is a matrix specifying the scores for observing particular amino acids at specific positions. In our study, this matrix is deduced from the training set. It has 4 rows and 21 columns, corresponding to 4 kinds of bases and 21 positions in the aligned ASs. When calculating the score for a new given AS sequence from the test set, we sum the negative log-odds score of that AS matching the motifs in PSSM. We use it to represent the similarity between a new test sample and the overall training samples.

###### Activity of siRNA oligonucleotides

It is affected by the secondary structure of the target mRNA, especially the target site [12] [13]. Unstructured siRNA duplexes can mediate more active silencing than those with base-paired ends, and the structure of the AS also affects potency [32]. We used RNAfold from the ViennaRNA package to predict the secondary structure of AS and mRNA sequences, including the minimum free energy (MFE), dot bracket notation (DBN), and centroid structure [33]. We used the MFE, the ratio of paired and unpaired bases in the DBN, as the features to describe the secondary structure of the AS and local mRNA. A smaller MFE means a higher affinity of targeted binding with more effective gene silencing. In addition, we took the secondary structure of the global mRNA into account to estimate the accessibility of the target sites more accurately.

###### GC content and GC stretch

GC content is the ratio of bases G and C in the AS. The GC stretch is the maximum length of continuous bases G or C. A relative study showed that the GC content in functional siRNAs has a trend of 30%-60% [34]. This might affect the potency via thermodynamic properties since the Gibbs energy between A and U is much larger than that between G and C. The GC stretch determines the lower bound of the overall stability.

#### 2.2.2 Convolution and Pooling Module

The predictor mainly uses a CNN to detect potentially decisive motifs for predicting inhibition. In the one-dimensional convolution and pooling module, there are 45 different kernels with multiple sizes elicited from the work [23]. These kernel sizes range from 5 to 20 for one-hot encoding of AS (M1), from 5 to 20 for RNA-FM encoding of AS (M2), and from 6 to 21 for one-hot encoding of the local mRNA (M5). The kernel in each convolution operation goes through all possible combinations of adjacent bases. We utilize the ReLU activation function to increase the nonlinear mapping capacity. Then, average pooling and maximum pooling follow each convolution operation. The purpose of pooling is to reduce the number of parameters, remove useless noises, and collect the representative patterns. For 45 different convolutions, this module produces a 90-dimensional vector (M7+M8+M9).

#### 2.2.3 Full Connection Module

In the fully connected module, we concatenate the pooling output (M7+M8+M9) and all other features together, including thermodynamic parameters (M3) and features extracted from expert knowledge (M4, M6). After batch normalization, the output is fed into four sequential submodules. Each submodule consists of one linear layer with ReLU activation and dropout. Finally, we use sigmoid activation to generate the prediction score, ranging from 0 to 1, the same as the inhibition label.

### 2.4 Experimental Setup

#### 2.4.1 Sampling and training

It is well known that machine learning-based algorithms face the problem of out-of-distribution (OOD) generalization. Therefore, we sorted the siRNA samples in an inhibition-descending order. During 10-fold cross-validation, we choose the samples with indices of i, i+10, i+20… (i=1, 2,…10) to form the test set to unify the distribution between the training set and test set. The Adam strategy is used to optimize the model with an initial learning rate set of 0.01. As a regression task, we train the model with the mean average error loss (MAE).

#### 2.4.2 Evaluation metrics

To assess the prediction capacity, we use two statistical metrics to evaluate our method: the Pearson correlation coefficient (PCC) and Spearman correlation coefficient (SPCC). PCC is used to evaluate the linear correlation between two sets of continuous values, while SPCC is used to assess the correlation of the discrete rankings. High SPCC is more meaningful in siRNA design than PCC. They can be calculated with the following equations:

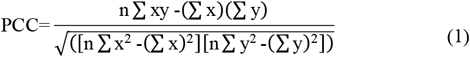

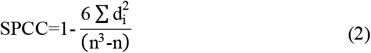

where x is the predicted value, y is the true value, n is the sample size, and di is the difference between the x rank and y rank for each pair of data.

## 3. Results and discussion

### 3.1 Comparison with other predictors

We compare our prediction model with the i-score, Biopredsi [17], DSIR [18], one CNN-based model [19], and one GNN-based model [35]. Due to unavailable webservers, we obtain the predictions of i-score, s-Bi-opredsi, and DSIR from the i-score website. Although some differences exist between s-Biopredsi and Biopredsi, the authors claimed that the correlation between them achieved a very high level of 0.997. In addition, we reimplement CNN-based work according to its description for the same reason.

We compare the performances of these models with 10-fold cross validation. The boxplot in **Fig. 3** shows that our predictor achieves the best performance, reaching an average PCC of 0.745 and SPCC of 0.736. The GNN-based model reached an average PCC of 0.725 and an SPCC of 0.7. The CNN baseline reached an average PCC of 0.671 and SPCC of 0.651, which are much lower than ours. The other three traditional models show unsatisfactory results.

**Fig. 3:**
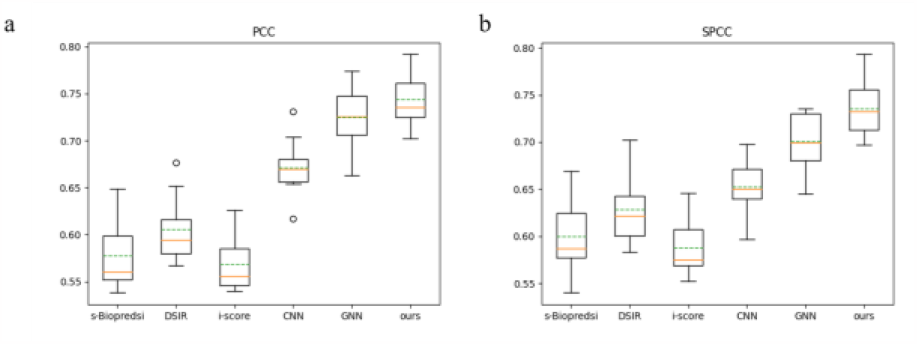
Comparison of siRNA silencing inhibition prediction of out model against other five tools on the same test set in 10-fold cross validation. Two metrics are: **a**. PCC, **b**. SPCC.

Our predictor yields better performance than the three traditional baseline methods. The reason may be that they mainly rely on manual feature engineering. However, our model utilizes a nonlinear mapping function in a wide and deep architecture to learn and select informative patterns. We can see that the performance of our predictor surpasses the CNN-based model, which merely extracts sequence features from local mRNA via one-hot encoding and thermodynamic parameters from AS. Other critical features are not used. In addition, the full connection module in the CNN-based model has only one linear layer with 25 nodes followed by a single node output. This structure is quite simple and unable to learn important information. It does not take any measure to avoid overfitting. On the two most important metrics in siRNA design, namely, PCC and SPCC, our model outperforms the GNN-based model. The reasons may be that it only takes the k-mer frequencies of siRNA and complete mRNA and thermodynamic properties as features. It ignores other significant features. Another serious problem is that this GNN-based model is trained and tested via deductive reasoning. That is, all siRNA samples from both the training set and test set are integrated into one graph. During training, only data from the training set are used to compute loss, while all weight matrixes are updated, including those for data from the test set. Therefore, the model cannot be used to predict new siRNAs that do not appear in fixed graph structures. Nonetheless, we can deduce that the biological mechanism of siRNA-mRNA interactions is quite suitable for modeling with graph structures to explore the inner deep relationship among different nodes.

In addition to the aforementioned five methods, we investigate a recently published method, pssRNAit, which utilizes SVM as the basic module. It integrates the inhibition scores evaluated by 12 different models, including DSIR, i-Score, Katoh, Reynolds, etc. It is trained and tested only on dataset-H by 5-fold cross validation, and it achieves an average PCC of 0.709 [36]. Considering that the web server of pssRNAit is also not accessible, we evaluate our model on dataset-H in the same way, achieving a better PCC of 0.738. This model aggregates different models’ predictions to strengthen its generalization instead of detecting new patterns.

### 3.2 Ablation study

To demonstrate the effectiveness of RNA-FM encoding and expert-knowledge-based features in our scheme, we perform ablation experiments in 10-fold cross validation. As shown in **Table 3**, our improvements produce meaningful results.

**Table 3.** The ablation study about RNF-FM encoding and expert-knowledge-based features.

| RNA-FM encoding | expert-knowledge-based features | average PCC | average SPCC |
| --- | --- | --- | --- |
| ✓ | ✓ | **0.745** | **0.736** |
| × | ✓ | 0.737 | 0.724 |
| ✓ | × | 0.728 | 0.719 |
| × | × | 0.718 | 0.708 |

The model achieves 1% average PCC and 1.1% average SPCC improvement when we only use the RNA-FM encoding method. It achieves a 1.9% average PCC and a 1.6% average SPCC improvement when we only use expert knowledge-based features. If both features are considered, our model will obtain the greatest enhancement. This indicates a significant 2.7% average PCC and 2.8% average SPCC improvement compared with the base model.

## 4. Conclusions

RNAi is a powerful technology used in posttranscriptional gene silencing. Multiple methods have been developed to predict and select effective siRNAs from numerous candidates. However, they are still not accurate and robust enough to design hyperfunctional siRNAs. In this study, we propose a novel deep-learning-based algorithm with multiple features as input, including siRNA and local target mRNA sequence context, thermodynamic properties, and other expert knowledge. Our model can automatically detect potential motifs from sequence embedding in a data-driven way instead of analyzing time- and labor-consuming biological experiments. These motifs may be potentially decisive in the actual inhibition of siRNA. We also use meaningful criteria from previous studies to further improve the accuracy. Experimental results show that our model outperforms all previous methods on the benchmark datasets.

In future works, incorporating chemical modification into siRNA design could be a potential direction. Some modifications can further improve the inhibition and repress off-target effects. Another possible direction is to utilize other algorithms, such as transformers, to replace the convolution, which may achieve a better result.

## 5 Declarations

### Ethics approval and consent to participate

Not applicable.

### Consent for publication

Not applicable.

### Availability of data and material

Refer to the supplementary files.

### Authors’ contributions

Bin Liu conducted the experiments and wrote the main manuscript text. Huiya Huang, Weixi Liao, Xiaoyong Pan, Cheng Jin, Ye Yuan reviewed the manuscript and gave instructions.

